# A multivariate analysis of CalEnviroScreen: comparing environmental and socioeconomic stressors versus chronic disease

**DOI:** 10.1101/221739

**Authors:** Ben K. Greenfield, Jayant Rajan, Thomas E. McKone

## Abstract

**Background:** The health-risk assessment paradigm is shifting from single stressor evaluation towards cumulative assessments of multiple stressors. Recent efforts to develop broad-scale public health hazard datasets provide an opportunity to develop and evaluate multiple exposure hazards in combination.

**Methods:** We performed a multivariate study of the spatial relationship between 12 indicators of environmental hazard, 5 indicators of socioeconomic hardship, and 3 health outcomes. Indicators were obtained from CalEnviroScreen (version 3.0), a publicly available environmental justice screening tool developed by the State of California Environmental Protection Agency. The indicators were compared to the total rate of hospitalization for 14 ICD-9 disease categories (a measure of disease burden) at the zip code tabulation area population level. We performed principal component analysis to visualize and reduce the CalEnviroScreen data and spatial autoregression to evaluate associations with disease burden.

**Results:** CalEnviroScreen was strongly associated with the first principal component (PC) from a principal component analysis (PCA) of all 20 variables (Spearman ρ = 0.95). In a PCA of the 12 environmental variables, two PC axes explained 43% of variance, with the first axis indicating industrial activity and air pollution, and the second associated with ground-level ozone, drinking water contamination and PM_2.5_. Mass of pesticides used in agriculture was poorly or negatively correlated with all other environmental indicators, and with the CalEnviroScreen calculation method, suggesting a limited ability of the method to capture agricultural exposures. In a PCA of the 5 socioeconomic variables, the first PC explained 66% of variance, representing overall socioeconomic hardship. In simultaneous autoregressive models, the first environmental and socioeconomic PCs were both significantly associated with the disease burden measure, but more model variation was explained by the socioeconomic PCs.

**Conclusions:** This study supports the use of CalEnviroScreen for its intended purpose of screening California regions for areas with high environmental exposure and population vulnerability. Study results further suggest a hypothesis that, compared to environmental pollutant exposure, socioeconomic status has greater impact on overall burden of disease.

## Background

A wide range of factors including demography, socioeconomic status, psychosocial stressors, and environmental exposures influence health outcomes [1–6]. One often used approach for addressing a single stressor is a risk assessment, which focuses on the probability of harm and is based on a quantitative convolution of exposure assessment with a dose-response assessment to provide an overall characterization of risk. But in order to protect vulnerable individuals and communities, environmental health science has broadened in emphasis from single-stressor evaluations to include integrated assessment of multiple stressors [1, 6–10]. This integration among disparate exposures presents a significant methodological challenge, requiring qualitative and less-formal quantitative methods that address hazard as the potential for harm [3, 4, 6, 7, 11, 12]. These impact assessments use metrics of exposure and dose-response but lack the quantitative direct link of these two factors that is common in risk assessment. Based on these methods, environmental justice advocates and health geographers have developed a variety of maps, indices, and tools that integrate environmental health hazards from multiple stressors at varying geographic scales. These tools incorporate a range of indicators including pollutant concentration or load estimates, contaminated sites or other hazards, built environment measures (e.g., urbanization, industry, green space, and road and traffic density), and population characteristics, such as educational attainment and socioeconomic status [2, 6, 8–10, 12–17]. A particular goal is the identification of geographic regions where vulnerable populations encounter environmental exposures, resulting in potential health impacts [2, 16, 18, 19]. To increase transparency and potential societal benefits, many integrated hazard assessment programs also engage the community at large in tool development and assessment, through community-based participatory research, solicitation of public commentary, and the provision and use of publicly accessible data [12, 13, 16, 20–22]. Some of these integration tools and methods also consider the potential interaction between environmental contributors to risk and the preexisting vulnerability of exposed populations to environmental stressors [1, 8, 16, 19, 23].

To complement the development of new methods, there is an ongoing need to quantitatively examine existing methods and tools. Critical and impartial evaluations will ensure that integrated assessment methods are well characterized, technically defensible, and appropriate for intended uses. Further, methods and their underlying data can be analyzed for geographic and statistical patterns of public health hazards [9, 10, 16, 23, 24]. The correlation among and between environmental exposures, socioeconomic vulnerability, and health outcomes also warrant investigation. In particular, an understanding of which health stressors (e.g., environmental, social, economic) are most associated with adverse health outcomes can aid in resource allocation and health policy direction across regions and populations [4]. For example, the relative health impact of environmental factors (e.g., pollution) versus population attributes (e.g., socioeconomic status, educational attainment) warrants examination. Multivariate methods (e.g., ordination, principal component analysis) and spatial statistics [25–27] can be very useful methods for these questions given the multivariate and spatial nature of health hazard assessment [9, 10].

An important case study for spatial health hazard evaluation is the California Communities Environmental Health Screening Tool version 3.0 (hereafter abbreviated as CalEnviroScreen). CalEnviroScreen was developed by the State of California Environmental Protection Agency’s (CalEPA) Office of Environmental Health Hazard Assessment (OEHHA) as a “screening methodology that can be used to help identify California communities that are disproportionately burdened by multiple sources of pollution [28].” CalEnviroScreen generates a numeric score, potentially ranging from 0 to 100. The score is based on 20 indicators: 12 measures of environmental exposure, 5 of socioeconomic vulnerability, and 3 of health outcomes (asthma, low birth weight, and cardiovascular disease) [13, 23, 24]. In addition to describing the spatial patterns of hazard, CalEnviroScreen is also intended to help guide state resource allocation. In particular, California Assembly Bill 32 (AB32) and Senate Bill 535 have established a cap and trade program for greenhouse gas emissions, and AB1550, passed in September 2016, requires that 25% of the anticipated one billion dollars of annual state revenue from this program be allocated to communities identified by CalEPA as having health vulnerabilities. CalEPA used CalEnviroScreen to identify these vulnerable communities [18, 23, 28].

The methodology for developing CalEnviroScreen has been detailed elsewhere, and CalEnviroScreen has previously been shown by the scientific development team to indicate strong racial disparities in environmental and socioeconomic vulnerability [23, 24]. The method and some of the underlying assumptions have also been subject to scrutiny as part of a public review process [21, 29], and a previous iteration (CalEnviroScreen version 2.0) was employed as a community disadvantage indicator, and found to be significantly associated with ovarian cancer survival [30]. Despite these applications and their policy significance and an October 10, 2017 search of all peer reviewed journal publications in Google Scholar and in PubMed that include the term “CalEnviroScreen” revealed no independent evaluation of CalEnviroScreen in the peer reviewed scientific literature. This is surprising, given that the CalEnviroScreen score will be used to decide on the allocation of around one billion dollars of state revenue, annually.

We examined the multivariate associations in the data underlying CalEnviroScreen (Version 3.0) and the statistical association between the 17 environmental and socioeconomic variables that describe environmental hazard and population vulnerability with an indicator of disease burden. We developed the disease burden indicator using publicly available hospital discharge data at the zip code scale.

## Motivation and Specific Aims

The literature on model performance evaluation makes clear the need for systematic efforts to show how a decision-support model performs relative to its intended objectives [31]. The work here was motivated by our observation that in the documentation to date for CalEnviroScreen, there is limited information regarding the multivariate associations in the variables used to construct the overall impact score. For example, there is no comparison to separate burden of disease measures, and no examination of the relationship between these associations and the CalEnviroScreen scores, themselves. Analyzing thirty sites at a zip code scale using a preliminary variable set and model formulation in 2012, Meehan August et al. [13] found correlations among many indicator variables proposed for inclusion in CalEnviroScreen, and low to moderate score sensitivity to changes in CalEnviroScreen model formulation. However, since that time the model formulation, variables, data sets, and geographic resolution have all changed. The current CalEnviroScreen model provides census tract scale data for over 8000 sites.

The CalEnviroScreen data reduction and score calculation methodology is moderately complex, and is based on considerable professional judgment [13, 23]. Public review comments frequently raise concerns that this calculation method may distort the comparative differences in hazard among locations, suggesting that comparison to an independently developed aggregate score would be useful. Therefore, one of our study objectives was to perform an external validation of the CalEnviroScreen score by comparison to a score obtained using simple application of a multivariate statistical algorithm (principal component analysis) to the underlying variables. We confront the need to quantitatively address outcome dependence on inputs in order to provide insight for decision makers.

In short, our aim was to systematically to describe associations among variables that go into the CalEnviroScreen score, whether the score strongly correlates with systematic change in the combined variables, and whether these indicators of hazard and vulnerability predict burden of disease. To achieve these objectives, we addressed four questions about the CalEnviroScreen source data and its use to produce an outcome score: 1. What are the correlation patterns and statistical associations in the variables? 2. How does the CalEnviroScreen numeric score compare to a score based on principal component analysis, employed as an alternate scoring method based on the statistical associations in the underlying data? 3. Using the alternate scoring method, what spatial patterns are evident in environmental hazard and population vulnerability in California? 4. Do the environmental hazard and population vulnerability indicators statistically predict overall disease burden? For this last question, we work to generate hypotheses regarding the ability of CalEnviroScreen to address health outcomes and the relative importance of environmental versus socioeconomic factors in determining disease burden.

## Methods

### CalEnviroScreen background and data

We downloaded the CalEnviroScreen data along with the CalEnviroScreen 3.0 scores as a Microsoft Excel spreadsheet file from the CalEnviroScreen website [32]. These data have been pre-cleaned and carefully prepared by CalEPA, as described elsewhere [23, 24]. The data set covers 8035 census tracts, though CalEnviroScreen 3.0 results are calculated for 7929 census tracts. CalEnviroScreen 3.0 includes 12 environmental hazard variables: ozone levels, concentrations of particulate matter ≤2.5 μm in diameter (hereafter, PM_2.5_), diesel particulate matter concentrations (diesel PM), traffic density, drinking water contamination, active pesticide mass used in agriculture (pesticide use), airborne toxic chemical releases, water body impairments, sites hazardous to groundwater (groundwater threats), sites targeted for cleanup, hazardous waste sites, and solid waste sites. CalEnviroScreen 3.0 also includes 5 socioeconomic vulnerability variables: low educational attainment, linguistic isolation, poverty, unemployment, and households severely burdened by housing costs (hereafter, housing burden). CalEnviroScreen 3.0 additionally includes three specific health outcome variables: asthma, low birth-weight, and cardiovascular disease, which are intended to indicate a combination of vulnerability to, and effects of, environmental exposures [23]. All variables were obtained based on data collected between 2009 and 2016, except drinking water contamination (2005 – 2013). Table S1 (Additional File 1) summarizes the variables, providing abbreviations, years represented, original units, and data transformations for this study. Cushing et al. [24], Faust et al. [23], and OEHHA [33] provide more extensive detail. The CalEnviroScreen 3.0 score is constructed from these data by a weighted averaging and multiplication algorithm, described elsewhere [23]. Briefly, each underlying variable is assigned a percentile rank score across each available census tract, a single weighted average is calculated for the 12 environmental variables and another weighted average for the 8 population variables (socioeconomic vulnerability and health outcome variables), each weighted average is linearly rescaled from 0 to 10, and finally, the two averages are multiplied achieving a score possibly ranging from 0 to 100.

### Disease burden measure

Although CalEnviroScreen does incorporate three specific health outcomes, we developed a separate disease burden measure independently of CalEnviroScreen to examine how well the environmental and socioeconomic variables in CalEnviroScreen predict overall burden of disease. To maintain transparency and potential for community access [4, 13], our disease burden indicator was developed using publicly available hospital discharge data collected at the zip-code level. Although some researchers may not consider hospitalization at zip code as the most specific measure that one would like for a disease burden analysis, it is what is publicly available to derive metrics of disease burden and has been used in other studies tracking links between environmental quality and disease [34, 35].

We characterized burden of diseases using discharge diagnostic codes (which used the ICD-9-CM schema). We obtained these data for all hospitalizations for a given calendar year using publicly available, de-identified, statewide hospital discharge data from the California Office of Statewide Health Planning and Development, spanning the years 2008-2011. Using these data, we classified hospitalizations by pre-determined ICD-9 diagnostic categories, focusing on diseases having environmental etiology. We determined the sum total number of hospitalizations for 14 disease diagnostic categories representing serious or chronic ailments known to have potential environmental etiology. The 14 categories were pneumonia, chronic obstructive pulmonary disease (COPD), asthma, myocardial infarction (MI), cerebrovascular accident (CVA), diarrhea, pancreatic cancer, lung cancer, breast cancer, lymphoma, leukemia, depression, schizophrenia, and low birth weight.

The overall approach of combining multiple diseases without weighting factors and assuming that hospital admissions can be a metric of regional disease patterns offers utility for impact assessment but has limitations. Combining multiple disease outcomes into a single metric of disease burden has the advantage of being comprehensive in capturing a large range of environmentally related diseases. The approach has been used in assessing global burden of disease for policy studies [36], but how or whether to use disease-weighting factors is a concern [37], and selection of weighting factors can both enhance and bias results. Because we were not interested in a metric of disability but only wide-scale disease patterns, we elected not to use disease-weighting factors. Using hospital admissions data as a metric of regional variations in disease patterns also has utility a health metric but limitations with regard to how well this metric relates to the health of a community. These issues have been addressed in the health tracking literature and show that the approach can be useful to policy makers [34, 38].

In a preliminary analysis, we found that the occurrence of these disease categories in hospitalized persons was positively correlated among the categories and also correlated with the total number of diagnoses (Additional File: Table S2, Fig. S1). Further, because the same patient hospitalization could have multiple diagnoses spanning multiple categories, rates for different disease categories are not mutually exclusive. To address these issues, and minimize the Type I error rate and complexity of the analysis, we assembled these data into a single count of total hospitalizations resulting from the 14 diagnostic categories. In this count, a hospitalization event that included more than one diagnostic category was recorded as a single event.

We summed the count of total hospitalizations at the zip code tabulation area (ZCTA) level, and divided by total population. For the denominator, we used ZCTA population estimates from the 2010 United States Census. We observed high variability in rates for populations below 100 individuals within a ZCTA (Additional File: Fig. S2). Therefore, we excluded ZCTAs having populations < 100 individuals from the final analysis to minimize the influence of statistical outliers.

In summary, this disease burden measure is the total rate of hospitalizations associated with at least one of the 14 diagnostic categories selected for a given ZCTA. Because the same person could in theory be admitted multiple times for the same diagnosis, the rates reported here are only approximations of population disease prevalence. They may be viewed as representing the impact of certain diseases (particularly chronic diseases), since apart from mortality, hospitalization is generally the most extreme result of any disease process.

Expecting chronic disease burden to be higher among the elderly, we obtained the percent of the population over 65 years old for each ZCTA from the 2010 US Census. This parameter (Over65) was used in addition to the environmental and socioeconomic variables derived from CalEnviroScreen as a predictor in statistical models used to explain the disease burden measure.

### Data preparation and spatial alignment

Hospitalization-related diagnoses were tabulated at the zip code scale, census population is at the ZCTA scale, and CalEnviroScreen data are available at the census-tract scale. Thus, spatial alignment was required prior to comparing CalEnviroScreen variables to the disease burden measure. The following data preprocessing steps were performed using ArcMap v10 (ESRI, Redlands, CA): 1. average the CalEnviroScreen data at the zip code scale to achieve a consistent analysis scale; 2. link the CalEnviroScreen and ICD-9 data together; 3. combine with a shapefile of zip code polygons; and 4. standardize the health outcome data to total population per zip code tabulation area (ZCTA). Additional File 1 further details this spatial alignment methodology.

### Statistical methods

Statistical analysis was performed in R (version 3.4.0) [39]. Principal component analysis (PCA) was conducted for data reduction, and to allow examination of the contribution to variance explained by individual variables (i.e., variable loadings) and the multivariate data structure, similar to other public health studies [10, 40, 41]. For all census tracts that had CalEnviroScreen results (n = 7929), PCA was performed on the correlation matrix using the R package factoMineR. Prior to PCA, missing values were imputed using the imputePCA command in the missMDA package [42].

Separate PCAs were performed to achieve different study objectives. A PCA was performed on all 20 CalEnviroScreen variables in combination in order to examine multivariate patterns of the entire data set. Two separate PCAs were also performed on the 12 environmental and the 5 socioeconomic variables to generate and evaluate a smaller number of variables representing the categories of environmental hazard, and socioeconomic status, respectively. Finally, another PCA was performed on the 17 environmental and socioeconomic variables. This PCA, which did not include the three health outcome variables (asthma, low birth-weight, or cardiovascular disease), was compared to the hospitalization rate disease burden measure (described above). The goal of this analysis was to examine how these exposure and population variables underlying CalEnviroScreen are generally associated with disease burden.

Environmental hazard and socioeconomic status variables (principal components) were compared to the disease burden measure (hospitalization rate for 14 diagnoses) using simultaneously autoregressive models (SAR), employing the R package spdep. SAR was chosen as a spatial autoregressive model appropriate to describe and test for linear relationships in the presence of spatial autocorrelation [27, 43]. Appropriate treatment of spatial autocorrelation was assessed based on Moran plots illustrating no association with spatially lagged means, global Moran’s I that was not significant, and a spatial dependence parameter (λ) that was significant via likelihood ratio test [43–45]. Models were selected based on minimizing the Bayesian Information Criterion (BIC). Parameter inclusion was based on reported p values (α = 0.05) and on ΔBIC, employing the rule of thumb that ΔBIC ≥ 2 provides positive evidence of model improvement [46]. Nagelkerke pseudo-R^2^ was calculated as a measure of model goodness of fit for SAR models. Analogous to traditional R^2^ in meaning (though not directly comparable), the Nagelkerke pseudo-R^2^ estimates from 0 to 1 the improvement in proportion of variation explained by the fitted model, versus a null (intercept-only) model [47]. In order to compare the contribution of each parameter to final variation explained by the model, the psuedo-R^2^ was compared between the full model and the model with that parameter removed.

Prior to statistical analysis, all variables were transformed to approximate a normal distribution and multivariate linearity required for linear model analysis [26, 48]. Transformations included log_10_ (7 variables), cube root (6 variables), square root (5 variables), and arcsine square root transformation (drinking water). PM_2.5_ and low birth weight did not require transformation (Additional File: Table S1). The combined disease burden measure (DB) exhibited skewness and long tails (leptokurtic) and standard transformations failed to achieve normally distributed model residuals. Normal residuals were therefore achieved employing a modulus transformation: 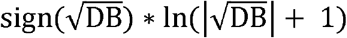 following John and Draper [49]. The predictor variables for the SAR were centered and scaled by subtracting the mean and dividing by the standard deviation. This converted the transformed variables (Additional File: Table S1) to the same unit normal distributions, such that a comparison of model coefficients would approximately indicate relative contribution of each variable to disease burden [50].

In the interest of independent assessment, we did not communicate with OEHHA, CalEPA, or any members of the CalEnviroScreen development team regarding any aspect of this study.

## Results

For Question 1, describing the multivariate structure of the CalEnviroScreen source data, we employ PCA to visualize which exposures are associated with each other and the prevailing patterns of overall exposure encountered in California. For Question 2, how CalEnviroScreen compares to an alternate metric based on PCA, we present the correlation between the main principal components and the CalEnviroScreen score. For Question 3, spatial patterns in environmental hazard and population vulnerability, we map the first principal component (PC) of separate PCAs performed on the environmental hazard and socioeconomic vulnerability variables. Finally, Question 4 examines and compares whether the exposure and vulnerability indicators in CalEnviroScreen predict disease burden. We employ the PCs rather than the individual parameters to focus on overall patterns of exposure and vulnerability and to reduce the number of required analyses. The relative importance of environmental versus socioeconomic parameters in the model illustrates which factors most influence disease burden.

### Multivariate data structure

Pearson’s pairwise correlation coefficients (r) indicate multiple associations for the underlying CalEnviroScreen data (Table 1). Positive pairwise associations are observed among variables related to air pollution and traffic, with diesel PM moderately correlated with PM_2.5_ (r = 0.41) and traffic density (r = 0.56), and toxic release correlated with these three variables (r = 0.43 – 0.56, Table 1). Socioeconomic indicators of vulnerability are also positively associated: low educational attainment, linguistic isolation, poverty, and unemployment exhibit r values ranging from 0.51 to 0.82, with the exception of linguistic isolation versus unemployment (r = 0.24) (Table 1; Additional File: Fig. S3). Housing burden is also positively related to these variables, with correlation coefficients ranging from r = 0.40 versus unemployment to r = 0.72 for poverty. The strongest correlation between socioeconomic and environmental variables is between linguistic isolation and diesel PM (r = 0.43). The strongest negative association among all variables is groundwater threats versus ozone (r = −0.33). Of the CalEnviroScreen health outcome variables, low birth weight is not well correlated with any environmental variables, but is highly correlated with housing burden (r = 0.72). Similarly, asthma is more positively correlated with the socioeconomic variables education (r = 0.51), poverty (0.53), and housing burden (0.50), than with any environmental variables. Cardiovascular disease exhibits moderate correlations with low educational attainment, poverty, and unemployment (r = 0.43 − 0.46) but also with the environmental variable, ozone (0.39).

**Table 1.** Correlation matrix of underlying data from CalEnviroScreen 3.0. Data were transformed and analyzed using pairwise Pearson’s correlation coefficients (r). Sample size ranged from 7694 to 8035 census tracts. Boldface indicates |r| ≥ 0.3.

|  | Ozone | PM<sub>2.5</sub> | Diesel PM | Traffic density | Drinking water | Pesticide use | Toxic release | Cleanup Sites | Groundwater threats | Hazardous waste sites | Water body impairments | Solid waste sites | Education | Linguistic isolation | Poverty | Unemployment | Housing burden | Asthma | Low birth rate |
| --- | --- | --- | --- | --- | --- | --- | --- | --- | --- | --- | --- | --- | --- | --- | --- | --- | --- | --- | --- |
| PM<sub>2.5</sub> | **0.41** |  |  |  |  |  |  |  |  |  |  |  |  |  |  |  |  |  |  |
| Diesel PM | −0.16 | **0.41** |  |  |  |  |  |  |  |  |  |  |  |  |  |  |  |  |  |
| Traffic density | −0.09 | 0.23 | **0.56** |  |  |  |  |  |  |  |  |  |  |  |  |  |  |  |  |
| Drinking water | **0.56** | **0.35** | −0.10 | −0.02 |  |  |  |  |  |  |  |  |  |  |  |  |  |  |  |
| Pesticide use | 0.08 | −0.05 | **−0.30** | **−0.30** | 0.21 |  |  |  |  |  |  |  |  |  |  |  |  |  |  |
| Toxic release | 0.06 | **0.56** | **0.54** | **0.43** | 0.09 | −0.24 |  |  |  |  |  |  |  |  |  |  |  |  |  |
| Cleanup sites | −0.12 | 0.10 | 0.20 | 0.14 | −0.01 | −0.01 | 0.13 |  |  |  |  |  |  |  |  |  |  |  |  |
| Groundwater threats | **−0.33** | −0.05 | 0.10 | 0.10 | −0.12 | 0.08 | 0.00 | **0.44** |  |  |  |  |  |  |  |  |  |  |  |
| Hazardous waste sites | −0.13 | 0.09 | 0.28 | 0.23 | −0.03 | −0.02 | 0.20 | **0.49** | **0.39** |  |  |  |  |  |  |  |  |  |  |
| Water body impairments | **−0.30** | −0.23 | −0.09 | −0.03 | −0.17 | 0.14 | −0.13 | 0.12 | 0.22 | 0.13 |  |  |  |  |  |  |  |  |  |
| Solid waste sites | 0.03 | −0.02 | −0.17 | −0.04 | 0.13 | 0.18 | −0.02 | **0.33** | 0.29 | **0.31** | 0.17 |  |  |  |  |  |  |  |  |
| Education | 0.06 | −0.10 | −0.18 | −0.08 | 0.06 | 0.09 | −0.12 | −0.08 | −0.03 | −0.13 | −0.12 | 0.19 |  |  |  |  |  |  |  |
| Linguistic isolation | −0.02 | **0.30** | **0.43** | 0.28 | 0.10 | −0.07 | 0.31 | 0.22 | 0.13 | 0.18 | −0.09 | 0.06 | **0.70** |  |  |  |  |  |  |
| Poverty | 0.23 | 0.22 | 0.16 | 0.05 | 0.18 | 0.04 | 0.04 | 0.19 | 0.13 | 0.12 | −0.11 | 0.17 | **0.82** | **0.58** |  |  |  |  |  |
| Unemployment | **0.30** | 0.16 | 0.00 | −0.05 | 0.19 | 0.04 | −0.01 | 0.05 | 0.02 | 0.02 | −0.08 | 0.10 | **0.51** | 0.24 | **0.60** |  |  |  |  |
| Housing burden | 0.04 | 0.15 | 0.29 | 0.26 | 0.06 | −0.16 | 0.17 | 0.15 | 0.10 | 0.14 | −0.08 | 0.08 | **0.57** | **0.50** | **0.72** | **0.40** |  |  |  |
| Asthma | 0.08 | 0.12 | 0.20 | 0.00 | −0.04 | −0.01 | 0.05 | 0.13 | 0.11 | 0.10 | −0.05 | 0.08 | **0.51** | 0.25 | **0.53** | **0.45** | **0.50** |  |  |
| Low birth weight | 0.10 | 0.18 | 0.19 | 0.11 | 0.06 | −0.09 | 0.14 | 0.08 | 0.00 | 0.09 | −0.09 | 0.03 | **0.33** | 0.25 | **0.32** | 0.27 | **0.72** | 0.28 |  |
| Cardiovascular disease | **0.39** | 0.16 | 0.02 | −0.08 | 0.18 | 0.08 | 0.03 | 0.03 | −0.05 | 0.01 | −0.14 | 0.08 | **0.46** | 0.17 | **0.43** | **0.44** | 0.21 | **0.71** | 0.24 |

### Principal component analysis and comparison to CalEnviroScreen

PCA was performed on the entire CalEnviroScreen data set (20 variables), on the data set without the health outcome variables (17 variables), and on the environmental (12 variables) and socioeconomic (5 variables) data. For the entire data set, the first three PCs explain 50% of data variability in combination. The first principal component (PC) (Fig. 1a, horizontal axis), explaining 23% of variation, is positively associated with all variables except for a weak negative association with pesticide use and impaired water bodies. The variables with the greatest variance along this axis are asthma (health indicator) and the five socioeconomic indicators (linguistic isolation, low educational attainment, poverty, unemployment, and housing burden). The other two health indicators (low birth weight and cardiovascular disease) are also associated with the first PC axis, as is PM_2.5_ and diesel PM.

**Fig. 1.**
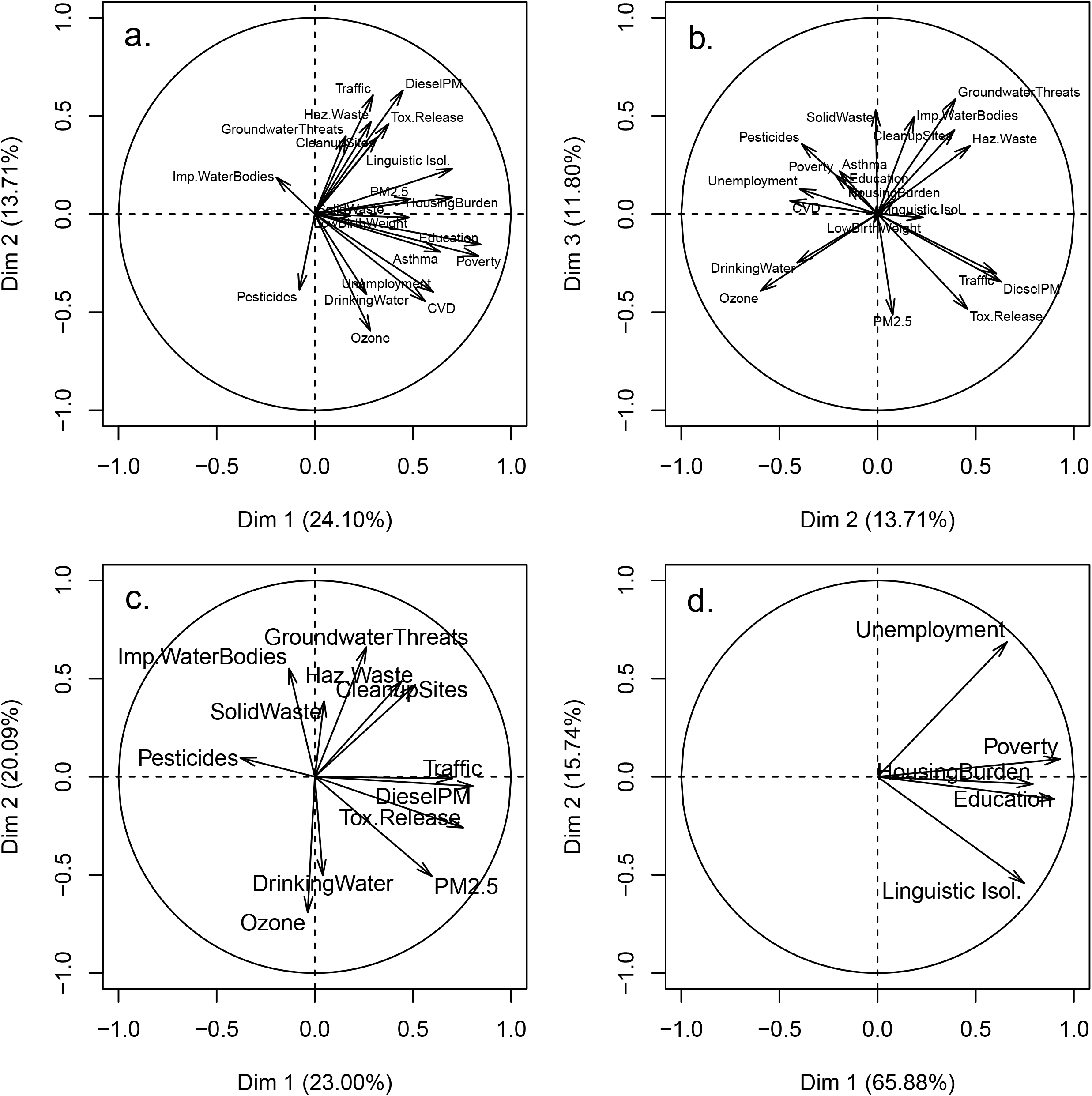
Results of principal component analysis of CalEnviroScreen environmental hazard and socioeconomic vulnerability variables across 7929 populated census tracts in California. Variability explained by individual principal components is in parentheses. a. All variables PC1 versus PC2. b. All variables PC2 versus PC3. c. Environmental variables only. d. Socioeconomic variables only.

The second PC (Fig. 1a, vertical axis; Fig. 1b, horizontal axis) explains 14% of variation. Indicators of industrial pollution and associated hazardous sites score positively with both the first PC (hereafter referred to as PC1all) and the second PC (hereafter, PC2all) (Fig. 1a, b). In particular, PC1all and PC2all are associated with groundwater threat sites, hazardous waste sites, cleanup sites, toxic release, traffic, and diesel PM. These variables are negatively correlated with pesticide use, which would be expected in rural areas, as well as drinking water contamination and ozone. Examining the biplot of PC2all and PC3all (Fig. 1b), we see a negative association between the polluted sites (cleanup sites, groundwater threats, impaired water bodies, and hazardous waste sites) and measures of drinking water contamination and ozone. Additionally, motor vehicle and industrial source-associated air pollution indicators (traffic, diesel PM, and toxic release) are negatively associated with pesticide use. Results for the PCA of all except the health outcome variables (17 remaining variables) are qualitatively very similar to that of the entire data set.

For the environmental PCA (Fig. 1c), the first two principal components explain 43% of data variability in combination. Along the axis of the first PC, explaining 23% of variance, there is a positive association among toxic releases and motor vehicle pollution indicators: PM2.5, diesel PM, and traffic. These are negatively associated with pesticide use. Sites contaminated due to industrial activity, including groundwater threats, hazardous waste sites, and cleanup sites, are positively associated with both the first and second PCs. Based on these associations, the first PC (hereafter, PC1env) represents general exposure to urban and industrial pollution. Interestingly, PC1env is strongly positively correlated at the zip code scale with population density (r = 0.81, n = 1602), indicating densely populated areas are more exposed to the main environmental hazards measured by CalEnviroScreen.

The second environmental PC (hereafter, PC2env) explains 20% of variance, and is negatively associated with ozone and drinking water contamination (Fig. 1c). Examination of the associations of PC2env indicates that elevated hazards due to ozone and drinking water contamination will tend to occur in different areas from impaired water bodies or groundwater threats.

For the socioeconomic PCA (Fig. 1d), the first two PCs explain 82% of data variance. The first PC (hereafter PC1soc) explains 66% of variance and is positively associated with all five socioeconomic indicators. Thus, PC1soc broadly indicates socioeconomic vulnerability. The second PC (PC2soc) explains 16% of variance, and separates unemployment from linguistic isolation (Fig. 1d).

The CalEnviroScreen method produces a numeric score calculated as a weighted sum of environmental variables (which CalEPA refers to as pollution burden) multiplied by a weighted sum of the socioeconomic and health outcome variables (referred to as population characteristics) [23, 24]. We were interested in how well this calculation method represents the prevailing multivariate patterns in environmental and socioeconomic vulnerability within the population of census tracts. That is, does CalEnviroScreen achieve its intended purpose of identifying areas exhibiting high hazard from a combination of environmental exposures and socioeconomic vulnerabilities? To evaluate the validity of CalEnviroScreen based on this criterion, we calculated Spearman’s rank correlation coefficient for the CalEnviroScreen score versus the main PCs from each PCA analysis, for all available census tracts (n = 7929). Spearman’s ρ was employed as a measure of association that is robust to nonlinear relationships, which were evident in this analysis (Fig. 2).

**Fig. 2.**
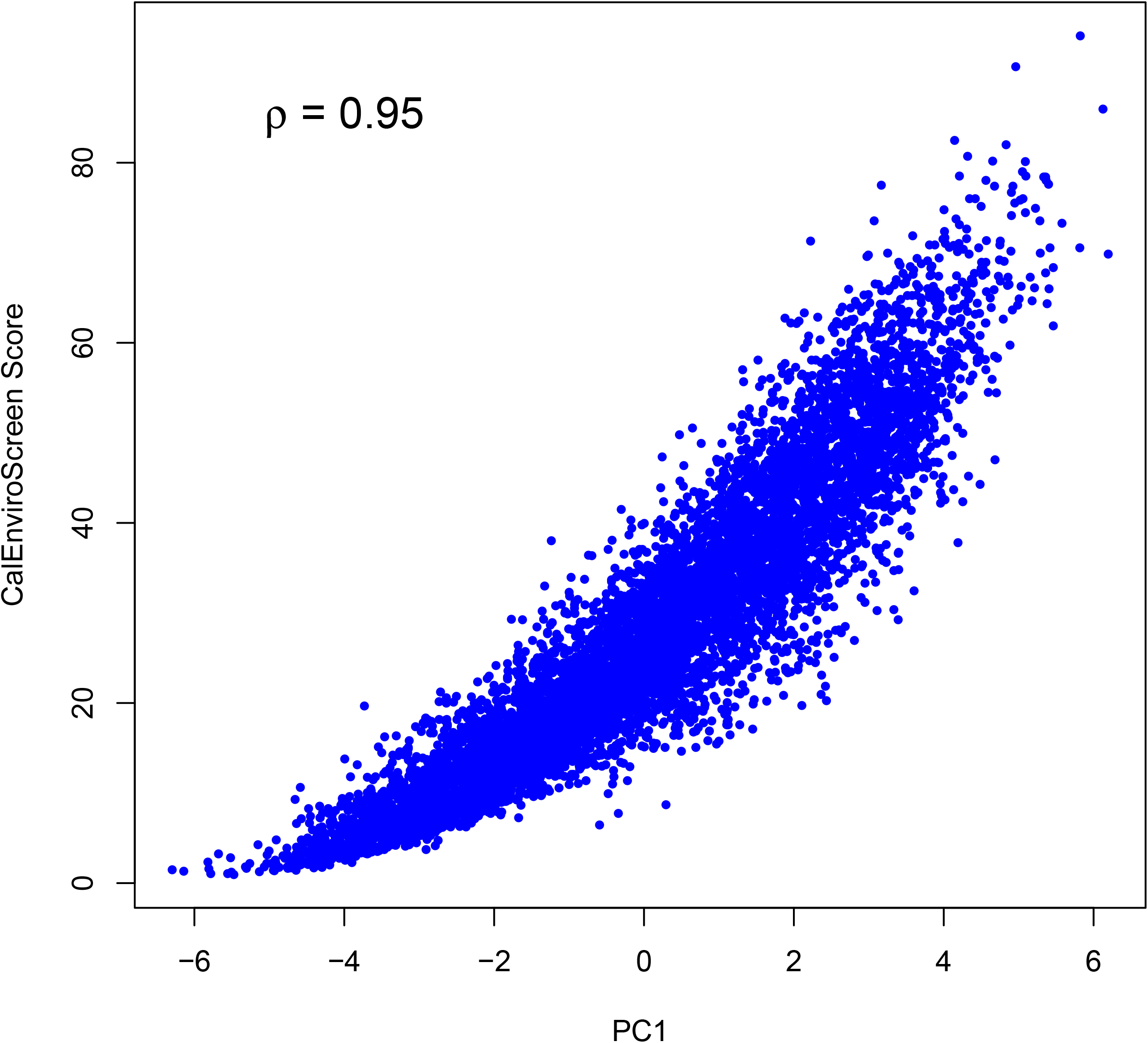
Association between first principal component for all variables (PC1) and CalEnviroScreen 3.0 score [23]. Each point represents a populated California census tract (n = 7929).

In the all-variables PCA, CalEnviroScreen is strongly associated with PC1all (ρ = 0.95; Fig. 2) and not associated with PC2all (ρ = 0.00) or PC3all (ρ = 0.10). When separate PCAs are performed on the environmental and socioeconomic data, CalEnviroScreen is strongly associated with PC1soc (ρ = 0.82), moderately associated with PC1env (ρ = 0.59), and not associated with PC2env (ρ = −0.08) or PC2soc (ρ = 0.00). These results indicate that changes in the predominant gradients underlying the data (PC1all, PC1soc, and PC1env) are generally captured by the CalEnviroScreen score. Thus, this single score effectively captures the prevailing gradients in the underlying variability in environmental exposures and socioeconomic vulnerabilities.

The only variables negatively associated with PC1all (Fig. 1a) were pesticides and impaired water bodies. Not surprisingly, the CalEnviroScreen 3.0 score exhibits no correlations with these two variables; Pearson’s r was 0.05 for pesticides and −0.04 for impaired water bodies. As a result, the CalEnviroScreen census tract rankings will be insensitive to these two variables.

### Spatial patterns

The spatial patterns in environmental hazard and population vulnerability can be seen by plotting the first principal component for the environmental and socioeconomic variables, respectively (Fig. 3, Fig. 4). For ease of visualization, we present these results aggregated at the zip code scale; spatial patterns in individual variables and calculated linear averages based on the CalEnviroScreen method are displayed at the census tract scale by the original authors [23]. Strikingly, both PC1Env and PC1Soc exhibit relatively high values in the southern Central Valley region of the state, especially the southwest San Joaquin Valley, indicating a pattern of elevated hazard and vulnerability in this region. PC1Env is also high in the more densely populated San Francisco Bay Area and Los Angeles regions, illustrating the abovementioned association between population density and the environmental hazard variables. In contrast to PC1Env, PC1Soc exhibits considerable heterogeneity in both the San Francisco and Los Angeles regions. All of these patterns were similarly observed using the CalEnviroScreen methodology for deriving their pollution burden and population characteristics metrics [23].

**Fig. 3.**
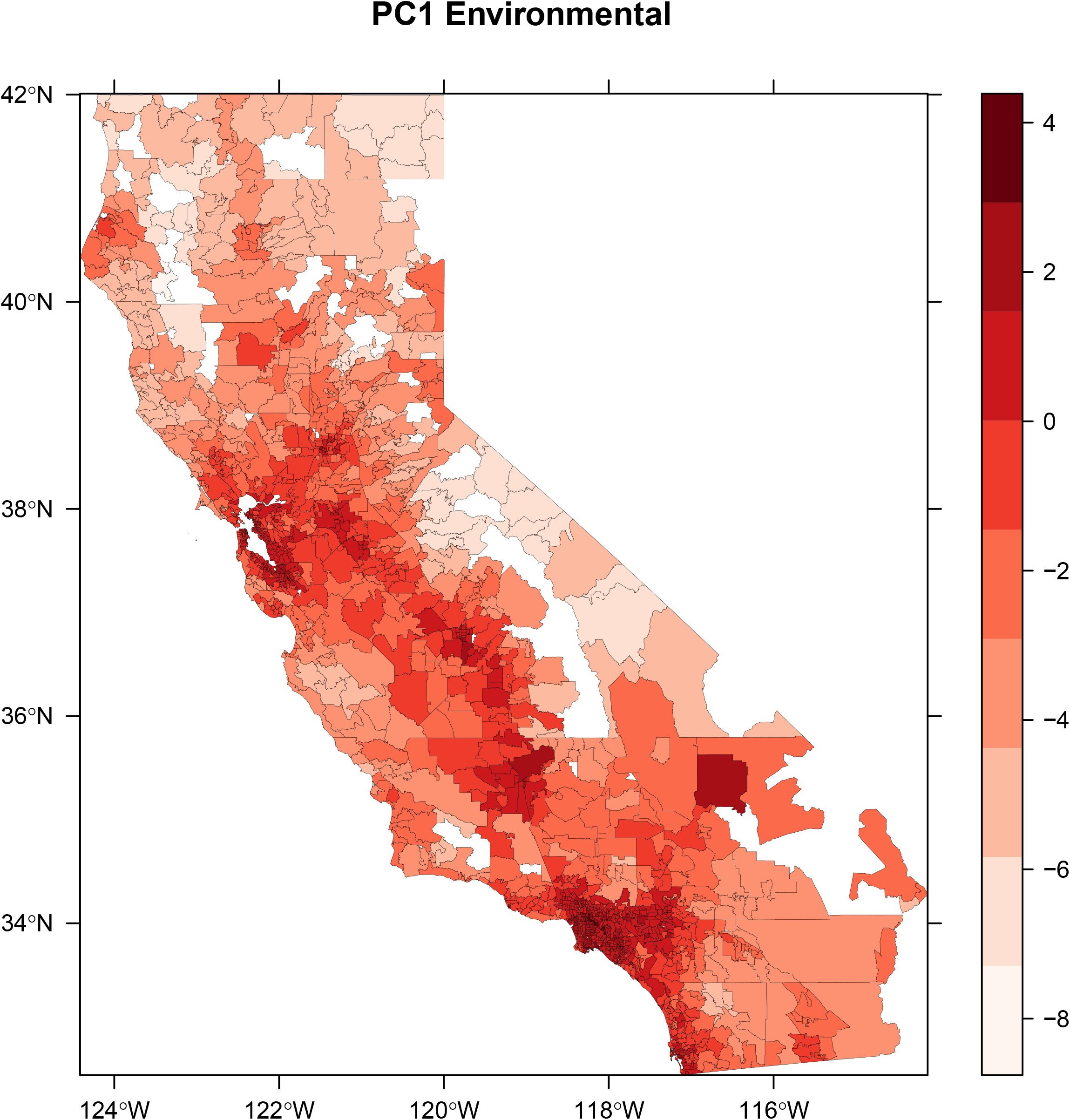
Spatial pattern in first principal component of environmental variables (PC1Env), aggregated at the zip code scale.

**Fig. 4.**
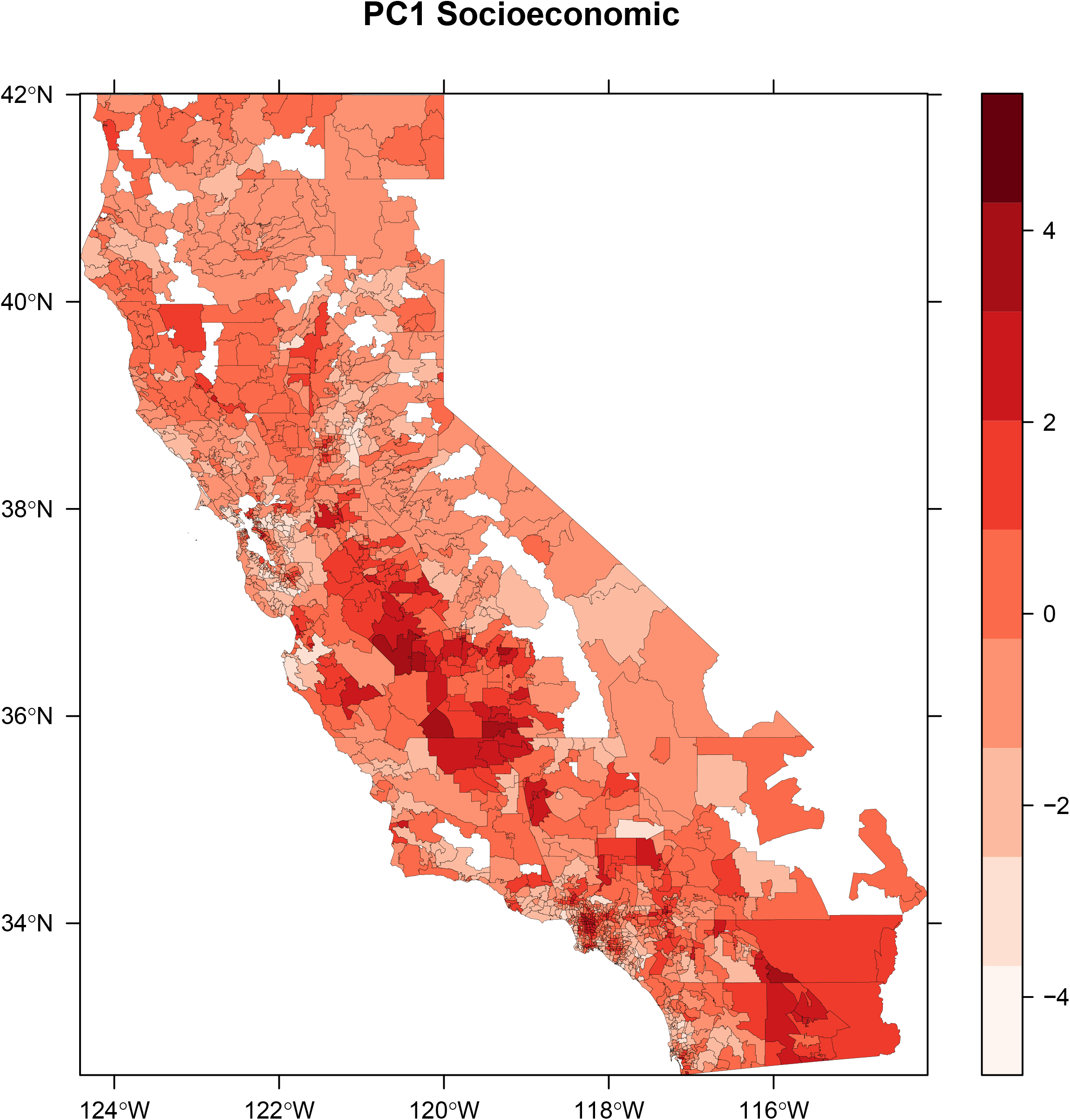
Spatial pattern in first principal component of socioeconomic variables (PC1Soc), zip code scale.

Spatial patterns in the CalEnviroScreen 3.0 measure (Fig. 5) clearly combine these two factors. As above (Fig. 2), the complex algorithm employed for aggregating the 20 variables in CalEnviroScreen [23] essentially captures the main underlying environmental and socioeconomic gradients (Fig. 3, Fig. 4). The highest scoring (and thus most impacted) areas are centered around the southwest San Joaquin Valley, peaking in the urbanized portions of Fresno County (including the cities of Fresno and Selma), as well as the Los Angeles region (Los Angeles, Pomona, and San Bernardino).

**Fig. 5.**
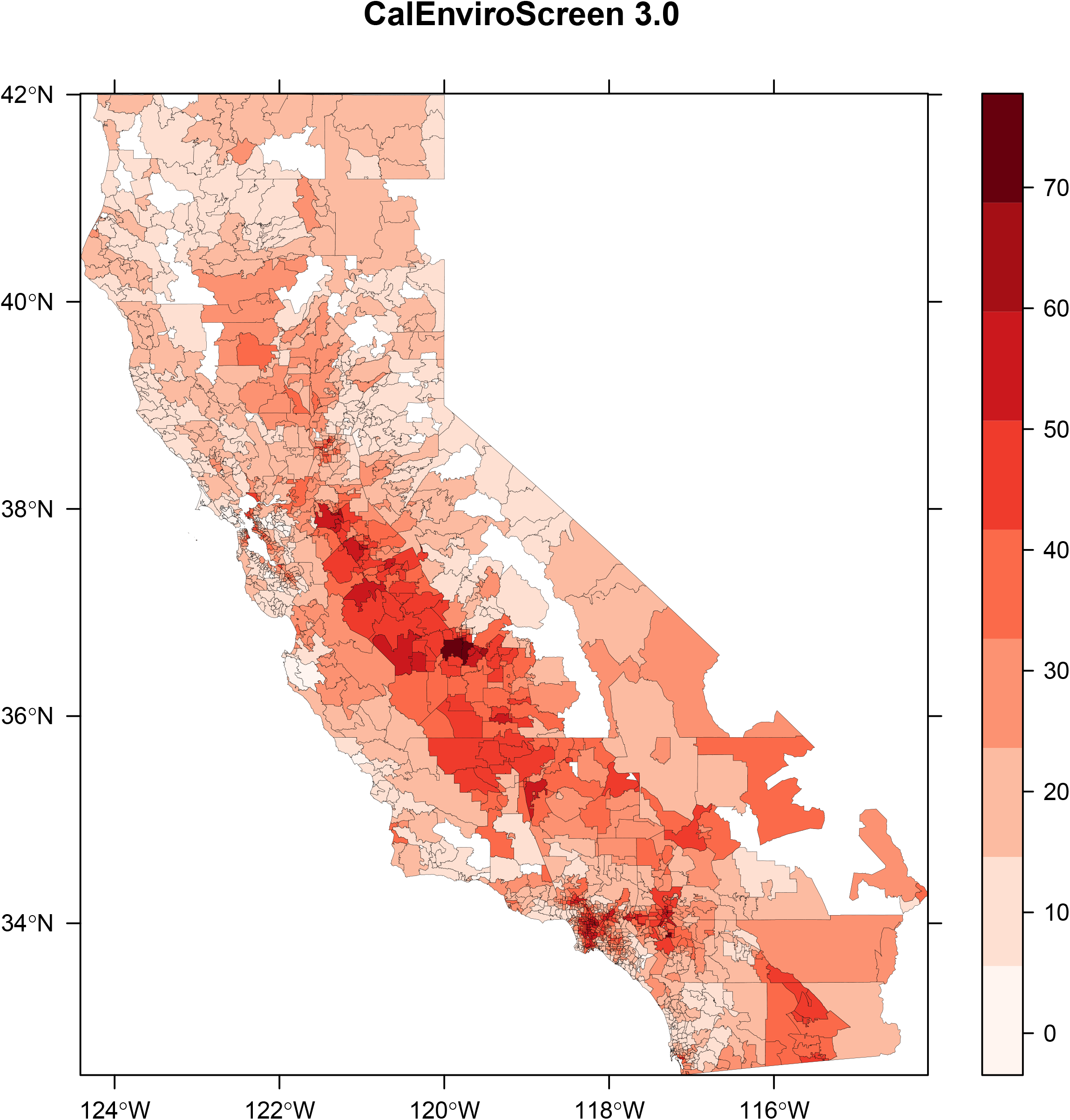
Spatial pattern in CalEnviroScreen 3.0, zip code scale. These data have been previously displayed at the census tract scale [23, 32], and are presented here to allow comparison to Figs. 3 and 4.

### Comparison to the disease burden measure

The principal components from the all environmental and socioeconomic variables PCA (PC1all, PC2all, and PC3all) and from the separated environmental and socioeconomic PCAs (PC1env, PC2env, PC1soc, and PC2soc) were evaluated as possible predictors for the disease burden measure that we developed. The predictor variables were not correlated with each other (|r| ≤ 0.13), with the exception of a weak negative correlation between PC1env and PC1soc (r = 0.31) and a weak correlation between PC1env and PC2soc (r = −0.31). Percent population > 65 years old (hereafter, Over65) was also included as a potential predictor. Over65 was moderately correlated with PC1all (r = −0.45), PC1soc (r = −0.36), and PC1env (r = −0.42), and uncorrelated with the other parameters.

Initial modeling using linear regression to predict disease burden indicated clear evidence of spatial autocorrelation of residuals for all models (based on Moran’s plots and significant Global Moran’s I). Thus, SAR was used to predict disease burden, and addressed these issues. For the SAR model on the all environmental and socioeconomic data PCA results (n = 1606 Zip Code Tabulation Areas), the following model form was obtained (coefficient estimate ± standard error, SE):

Disease burden = (0.47 ± 0.031)PC1all + (0.59 ± 0.022)Over65

For both PC1all and Over65, ΔBIC > 2, indicating that addition of each parameter was important in explaining disease burden, and p < 0.0001. In contrast, PC2all (p = 0.10), PC3all (p = 0.96), and the intercept (p = 0.97) were not significant as added model terms. For the final model, the Nagelkerke pseudo-R^2^ (hereafter, R^2^) was 0.49, which was effectively unchanged when attempting to include PC2all or PC3all. Change in R^2^ after removing individual parameters illustrates the contribution of each parameter to explaining the variability in disease burden. Decrease was greater when removing Over65 (new R^2^ = 0.25) than when removing PC1all (new R^2^ = 0.42). The association with PC1all generally describes the association of disease burden with the correlated variability in all of the CalEnviroScreen environmental and socioeconomic indicators except for pesticide use and impaired water bodies (Fig. 1a).

For the separate environmental and socioeconomic PCA results (N = 1606), three model terms contributed to describing disease burden (coefficient estimate ± SE):

Disease Burden = (0.35 ± 0.035)PC1env + (0.31 ± 0.024)PC1soc + (0.21 ± 0.026)PC2soc + (0.60 ± 0.021)Over65

ΔBIC > 2 and individual parameter p-values were < 0.001 for all included model terms. Neither PC2env (p = 0.16) nor the intercept (p = 0.93) were significant. The model coefficient was largest for Over65, simply indicating that age must be accounted for in the analysis. The positive coefficient for PC1env and PC1soc indicate an overall association with environmental exposures and socioeconomic vulnerability. The coefficient for PC2soc further suggests an association with unemployment, rather than linguistic isolation, in particular. The association with two socioeconomic principal components but just one environmental principal component, with qualitatively similar coefficient magnitudes on standardized variables, suggest a greater importance of socioeconomic than environmental factors in predicting overall disease burden. Similarly, the Nagelkerke pseudo-R^2^ indicates more variability explained by socioeconomic variables. Compared to the full model (R^2^ = 0.51), the R^2^ slightly declined when removing PC1env (R^2^ = 0.48), PC1soc (R^2^ = 0.46), or PC2soc (R^2^ = 0.49) from the model, but declined more when removing Over65 (R^2^ = 0.27), or PC1soc and PC2soc in combination (R^2^ = 0.44). These results suggest that whereas environmental hazard and socioeconomic vulnerability both contribute, socioeconomic vulnerability is more important than environmental hazard for explaining disease burden at the zip code scale.

## Discussion

### Multivariate analysis of the California Communities Environmental Health Screening Tool

Our study results support the use of CalEnviroScreen as a census tract scale indicator of environmental health hazard and vulnerability. The CalEnviroScreen numeric score was strongly associated with the first principal components in all analyses, indicating that it represents the primary underlying gradients within the data set. Additionally, the spatial patterns in the first principal components matched those of the CalEnviroScreen combined indicators and numeric score. Further, the principal components from the CalEnviroScreen exposure and vulnerability variables were significantly associated with our estimate of disease burden, which is a general indicator of health care burden. Models that contained these variables, and also accounted for spatial autocorrelation and proportion of population that was over 65, explained approximately 50% of the variation in the underlying data. Given all of the uncertainty and assumptions with the study data and scale, this suggests that the CalEnviroScreen data includes multiple exposure hazards and socioeconomic vulnerability indicators, which in combination influence burden of disease. Our analysis, therefore, suggests that CalEnviroScreen is an appropriate tool for its intended purpose: to identify vulnerable communities for resource allocation in environmental health restoration [18].

The first few principal components explained limited variance in the underlying data set, and many of the parameters, especially environmental exposure measures, were weakly correlated. Similar to our results, Lalloué et al. [9] observed 55% of total variance explained in the first three components of a multiple factor analysis of 17 environmental indicators and Messer et al. [10] observed 23 to 46% of variability explained in the first component in five related domains (air, water, land, built, and sociodemographic). These similar observations of low to moderate variance explained in the first principal components suggest that it is not possible to explain much variability in environmental health hazard using a small subset of indicators. Rather, a complete accounting of environmental hazards and population vulnerability requires a wide range of indicators, as employed in CalEnviroScreen and other studies [9, 10, 12–14, 23]. The limited variability explained by the first few principal components further suggests that for the 20 hazard parameters captured in CalEnviroScreen 3.0, there will be many exposure, vulnerability, and health outcome combinations that are not fully described by combined multivariate gradients. For example, the negative association of ozone air pollution with groundwater threats and water body impairments is not readily explained but suggests that residents of different regions encounter different exposure hazards. Further examination of the statistical properties and demographic vulnerability of sites exhibiting unique exposure combinations is ongoing. These efforts, performed by local agencies, interest groups, and community-based organizations in evaluating updates to the CalEnviroScreen method [51], complement the CalEnviroScreen numeric score by establishing a typology of vulnerable communities. How to incorporate these efforts into resource allocation decisions remains a difficult policy challenge.

The evaluation and modification of CalEnviroScreen is reflected in the recent release of CalEnviroScreen 3.0, updating much of the data, and attempting to address prior community review comments. In comparison to CalEnviroScreen 2.0, CalEnviroScreen 3.0 added two new indicators (hospital visits for heart attack, and low income households burdened by high housing costs), removed a vulnerable age indicator, and retained the other 18 indicators included in this paper [23]. Despite these changes, the multivariate patterns we observed in the correlation coefficients and principal component analysis results were almost identical between the two versions, and we therefore chose to focus on CalEnviroScreen 3.0 in this paper. Continued examination of the statistical properties and association with health outcomes is warranted for this and other multivariate hazard screening methods [9, 16, 17, 40], to complement the ongoing public discourse and review.

Some policy interventions may best be geographically targeted using additional information beyond the CalEnviroScreen score itself. Most of the environmental hazards measured were associated with urbanization and industrial activities, and the first environmental PC was strongly associated with population density. Although environmental exposure hazards are often elevated in urban areas [10], this is not always the case. For example, agricultural pesticide exposure is linked to a variety of developmental and health effects [52–54]. The pesticide use indicator in the CalEnviroScreen metric largely derived from agricultural application [23], was uncorrelated or negatively associated with the other environmental hazard indicators, and uncorrelated with the metric itself. Elevated nitrate concentrations in drinking water, another agricultural pollutant, is an additional known hazard for Latino communities in the San Joaquin Valley [55]. Given that rural communities tend to have lower incomes and reduced access to medical care [10, 56], additional measures of exposure hazard and vulnerability that affect isolated populations may warrant consideration alongside the CalEnviroScreen score.

### Disease burden was more associated with socioeconomic status than environmental hazards

We employed multivariable analytical methods to separate out chemical pollutant exposure hazard versus socioeconomic variation within California. Although environmental hazards and socioeconomic vulnerabilities are often correlated through complex causal pathways, and low income communities often face disproportionate environmental exposures [17], our multivariate approach allowed for a direct comparison of the statistical effect of environmental hazard versus socioeconomic vulnerability indicators on burden of disease. This is because the principal components for environmental variation (PC1env, PC2env) and for socioeconomic status (PC1soc, PC2soc) were uncorrelated with each other.

The association of the disease burden variable with both socioeconomic principal components but only one environmental principal component, as well as the greater combined contribution to variability explained (change in R^2^) suggest a stronger association of disease burden with socioeconomic status than with environmental pollution exposure. This was also evident in the greater correlation of individual socioeconomic variables with cardiovascular disease, asthma, and low birth weight. This supports the paradigm that underlying population vulnerability, resulting from socioeconomic conditions, must be considered in health risk assessment [1, 3, 12]. This finding is further acknowledged in that investigations of the environmental causes of disease typically adjust for indicators of socioeconomic status [5, 17]. The generalizability of our finding that socioeconomic factors better explained disease burden than environmental hazards merits investigation, as it would have implications for intervention priorities, as well as for conceptualization of the primary structuring factors that influence disease.

The multivariate and exploratory approach of our study reflects objectives quite different from a traditional epidemiological evaluation of how one or a small number of exposures affects a single outcome. Instead, our approach falls within the realm of quantitative methods for comparing among and evaluating cumulative environmental impacts in combination [6, 9, 10, 17]. We identified prevailing gradients of exposure and vulnerability, and observed how these patterns were associated with disease burden. We observed relatively strong associations among all of the socioeconomic indicators (education, income, unemployment, linguistic isolation), each of which may exhibit a separate impact on vulnerability [13]. This could explain the stronger association between socioeconomic indicators and disease burden, in contrast to environmental hazards, which were less correlated, such that the gradients in multivariate exposures were weaker. In other words, our findings support the paradigm that population disease burden will be more strongly impacted when multiple stressors occur in combination. As such, examination of the multivariate association among stressors should provide added and complementary information to bivariate analyses of exposure versus outcome.

### Limitations and caveats

Like all census-tract-scale studies of publicly available spatial exposure and health data, this study has limitations. For their similar study of the San Joaquin Valley region, Huang and London [14] eloquently describe the limitations of studies using publicly-available spatial exposure data. Our study does not establish causality and we cannot extrapolate inferences to the individual level [57]. We used publicly available health outcome data; as such our analysis was restricted to hospital discharge data at the zip code scale, which can be used to indicate overall morbidity [38], but may also be subject to bias [58]. Moreover, the specific choices we made regarding ICD-9 endpoints that had environmental etiology could be questioned, and must be interpreted as a general burden of disease measure, rather than indicative of any specific health outcome. Data required geographic alignment, including assembly of different parameters provided at multiple and varying spatial scales. In particular, CalEnviroScreen data were available at the census-tract level, the disease burden measure at the USPS zip code level, and spatial polygon arrangement at the zip code tabulation area-level. Inaccuracies are inevitably introduced when aligning these different spatial scales [59]. In line with the protection of individual rights to anonymity in publicly accessible outcome data, individual-level demographic information was masked, and residential addresses were limited to USPS zip code. These factors likely in part explain the limited strength of associations observed in this study. In particular, the association between the environmental principal components and disease burden was weak in our study. This was similar to Gaffron and Niemeier [19], who observed a very low bivariate strength of association (R^2^ = 0.018) between PM_2.5_ (environmental variable) and emergency visits for asthma (health outcome) employing CalEnviroScreen census tract data in six Sacramento, CA region counties. Especially, studies are needed combining longitudinal data sets of disease occurrence with CalEnviroScreen and other hazard measures [30].

## Conclusions

Previous studies at similar spatial scales and resolutions have established relationships of environmental hazards and disease risk with race and socioeconomic status, with implications for resource allocation and policy [14, 15, 19, 24, 60]. Prior studies have also shown geographic indicators of socioeconomic status or vulnerability to be associated with hospitalization rates or disease occurrence [2, 30]. However, few studies explicitly evaluate and describe the multiple patterns of association that occur across a range of health hazards and vulnerabilities at the census-tract scale [9, 10]. We use this approach to evaluate a cumulative impact screening methodology (CalEnviroScreen), observing that the methodology produced a spatial data set that captures the main underlying gradients, which in turn are associated with the burden of diseases having environmental etiology. CalEnviroScreen should therefore be useful for its intended purpose of screening for those communities most vulnerable to environmental exposure.

We observed that socioeconomic indicators were associated with each other and contributed to explaining disease burden, and that an environmental gradient of urban and industrial pollution also contributed to explaining disease burden. Ground-level ozone and drinking water threats were negatively associated with impaired water bodies and groundwater threats. Some of these findings corroborate findings from the preliminary analysis for CalEnviroScreen development using 30 zip codes by Meehan August et al. [13]. The existence of separate gradients of environmental hazard and socioeconomic disparity, and the varying ability to predict disease burden highlight the need for continued emphasis on integrated approaches in vulnerability assessment.

## Additional files

**Additional file 1:** Supporting information (PDF, 11 pages). Includes the following: methods for spatial joining of CalEnviroScreen versus hospital data; correlation of variables in ICD-9 hospital data (text and Table S2); study variables description (Table S1); scatterplot matrix of hospital visit rate parameters (Fig. S1); total hospital ICD-9 codes vs. total population (Fig. S2); and scatterplot matrix of socioeconomic variables (Fig. S3).

## Abbreviations

BIC: Bayesian Information Criterion
California Communities Environmental Health Screening Tool Version 2.0: CalEnviroScreen
CalEPA: California Environmental Protection Agency
COPD: Chronic obstructive pulmonary disease
CVA: Cerebrovascular accident
DB: Disease burden measure
ICD-9-CM: International Classification of Diseases 633 Ninth Revision Clinical Modification
MI: Myocardial infarction
PC: Principal component
PCA: Principal component analysis
PM: Particulate matter
PM_2.5_: Particulate matter 2.5 μm in diameter or below
r: Pearson’s pairwise correlation coefficient
SAR: Simultaneously autoregressive model
ZCTA: Zip code tabulation area

## Declarations

### Ethics approval and consent to participate

Not applicable. This study analyzes only existing, publicly available, de-identified data, analyzed at an aggregated scale.

### Consent for publication

Not applicable

### Availability of data and materials

The study was performed using publicly available data as described in the text. The datasets used and analyzed during the current study are available from the original sources or from the corresponding author on reasonable request.

### Authors’ contributions

BG and JR conceived the study, developed study methods, and prepared the study data. BG performed the analyses and wrote the manuscript. JR and TM reviewed the paper and provided comments and edits. TM and BG addressed reviewer comments and revised the manuscript. All authors have read and approved the manuscript.

## Acknowledgements

The data for CalEnviroScreen, which forms the basis of this analysis, has been made publicly available by California EPA’s Office of Environmental Health Hazard Assessment. We thank Gregory S. Biging and Alejandra Benitez for helpful statistical feedback and John Balmes for constructively reviewing manuscript drafts. We thank Lynne Messer and Martin Lawrence for reviewing the manuscript.

